# Rapid withdrawal from a threatening animal is movement-specific and mediated by reflex-like neural processing

**DOI:** 10.1101/2023.01.29.526084

**Authors:** Henry Railo, Nelli Kraufvelin, Jussi Santalahti, Teemu Laine

## Abstract

Responses to potentially dangerous stimuli are among the most basic animal behaviors. While research has shown that threats automatically capture the attention of human participants, research has failed to demonstrate automatic behavioral responses to threats in humans. Using a novel naturalistic paradigm, we show that two species of animals humans often report fearing trigger rapid withdrawal responses: participants withdrew their arm from photos of snakes and spiders faster, and with higher acceleration when compared to bird and butterfly stimuli. The behavior was specific to withdrawal as approach movements or button-press/release tasks failed to detect a similar difference. Moreover, between-participant differences in how aversive they found the stimuli predicted the participant’s withdrawal speed, indicating that the paradigm was also sensitive to trait-level differences between individuals. Using electroencephalography (EEG), we show that the fast withdrawal was mediated by two attentional processes. First, fast withdrawal responses were associated with early amplification of sensory signals (40-110 ms after stimulus). Second, a later correlate of feature-based attention (early posterior negativity, EPN, 200-240 ms after stimulus) revealed the opposite pattern: Stronger EPN was associated with slower behavioral responses, suggesting that the deployment of attention towards the threatening stimulus features, or failure to “disengage” attention from the stimulus, was detrimental for withdrawal speed. Altogether, the results suggest that rapid behavioral withdrawal from a threatening animal is mediated by reflex-like attentional processing, and later, conscious attention to stimulus features may hinder escaping the treat.

## Introduction

Humans are assumed to be equipped with neural circuits that enable them to automatically detect, and consequently avoid potentially dangerous stimuli (henceforth, “threat-relevant stimuli”) (LeDoux, 2012; Öhman & Mineka, 2001). Snakes have been proposed as prototypical threat-relevant stimuli that humans typically avoid and fear due to automatically activated neural circuits. However, despite the common anecdotal narrative that humans reflexively jump away from snakes when they encounter them in the wild, scientific evidence for the claim that snakes, or other types of threatening stimuli trigger automatic avoidance behaviors is lacking. Here, using a novel “naturalistic” experimental approach where participants interact with visual stimuli with their hand, we investigated behavioral avoidance reactions, and their electrophysiological correlates in human participants.

Snake detection theory states that primates evolved an automatic capacity to detect snakes because snakes were the main predators of early primates (Isbell, 2019; Öhman & Mineka, 2003). This phylogenetic adaptation is assumed to also be present in humans: a primitive neural pathway that relays information from the retina to amygdala via pulvinar is assumed to signal the presence of threats, and activate behavioral responses to the threat (LeDoux, 2012; Öhman & Mineka, 2001). Indeed, research on monkeys shows that the pulvinar responds rapidly and strongly to pictures of snakes (Le et al., 2013, 2016), and consistent with the proposed evolutionary basis of snake detection, primates seem to be innately “prepared” to develop change-resistent fear towards animals such as snakes and spiders (Kawai & Koda, 2016; Öhman & Mineka, 2001; Seligman, 1971). Other studies suggest that in addition to subcortical “threat detection pathways”, threat detection also depends on cortical areas that enable participants to learn to detect threat-relevant stimuli, and determine the adequate behavioral response to it depending on the context (Li & Keil, 2023; Lobue & Adolph, 2019; Pessoa, 2010; Pessoa & Adolphs, 2010).

In humans, event-related potentials (ERP) have been widely employed to characterize the time-course of activation that leads to the detection of a threat. Fear conditioning studies have reported amplified ERP responses in the C1 time window (peak around 50 ms), suggesting that already the earliest wave of visual stimulus evoked activity may be sensitive to threat-related information (Hintze et al., 2014; Sperl et al., 2021; Stolarova et al., 2006; Thigpen et al., 2017). The following P1 ERP wave (peaking around 75 ms after stimulus onset) is sensitive to unpleasant valence images, suggesting that stimuli such as potentially threatening objects receive priority early on in visual processing (Carretié et al., 2006; Olofsson et al., 2008a; Smith et al., 2003). The established neural correlate of snake detection in humans is the Early Posterior Negativity (EPN) taking place later in visual processing (around 200 ms) (Grassini et al., 2019; He et al., 2014; van Strien et al., 2014; Van Strien et al., 2016). According to the prevailing interpretation, the EPN reflects the deployment of feature-based selective attention towards the potentially threatening visual object (Dolan, 2002; Pessoa, 2010; Schupp et al., 2004, 2007). The (often implicit) assumption in many previous studies is that threat-evoked changes in attentional processing translate to behavioral responding: When a stimulus such as snake captures the participants attention, it also leads to improved behavioral responses (e.g., faster escape from the threat). However, crucially, previous studies typically examine ERP correlates of threat-processing in isolation, without also examining concurrent changes in behavior.

Behavioral evidence suggests that when humans perform a visual search task, threat-relevant stimuli such as snakes automatically capture their attention: Participants are faster to detect a threat-relevant target (such as a snake) among control images, and slower to find the non-threatening target when threat-relevant distractors are also present (March et al., 2017; Öhman et al., 2001; Öhman & Mineka, 2001; Soares et al., 2014; Zsidó et al., 2023). Yet, reaction times in the visual search paradigm are often long, meaning that they fail to provide evidence that treat-relevant stimuli evoke rapid behavioral responses such as withdrawal. The visual search task literature has also been criticized for methodological problems (Quinlan, 2013). Threat-detection has also been studied with paradigms such as the dot-probe tasks, but the majority of these studies have failed to find evidence for automatic allocation of attention towards threatening stimuli (MacNamara et al., 2013). Previous studies have mostly examined human participants’ responses in laboratory reaction time tasks, but very little is known about participants’ behavior in more ecologically valid situations. On exception to this is the study by Rinck et al. (2021) which showed that when participants interacted with stimuli on a touch-screen monitor, participants were slower to grab spider images (when compared to leaf stimuli). In another study, Rinck et al. (2010) applying an immersive virtual reality environment, showed that participants who are fearful of spiders spent more time looking at spiders when compared to non-fearful participants.

We reasoned that if humans are equipped with phylogenetic neural adaptations that mediate behavioral responses to threat-relevant stimuli, these neural processes may trigger specific, stereotypical responses such as arm withdrawal rather than speeding up all types of movements. If this is the case, the behavioral response may need to be measured using a naturalistic paradigm instead of, for instance, a button-press. To this end, we developed a paradigm where a participant directly interacts with a stimulus presented on a large touchscreen monitor by either touching it (i.e., approach movement), or withdrawing from it (Fig. 1a). The participants performed a go/no-go task where their task was to respond (either by approach or withdrawal movement) if the image contained an animal (snake, spider, bird, or a butterfly). When the image did not contain an animal (these images included pine cones, mushrooms, leaves, or flowers), the participant was required to keep their hand still.

**Figure 1.**
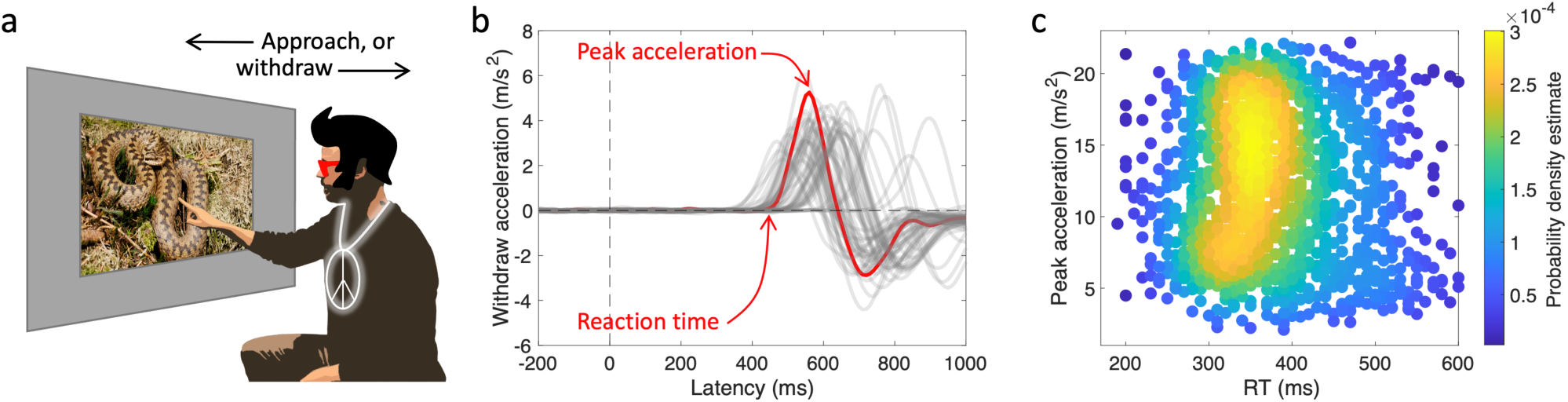
Behavioral paradigm and measures. a) Participant’s task was to either withdraw from, or approach with his/her hand an animal presented on a large touch screen monitor. The participant was instructed to not to move his/her hand when the image did not contain an animal (i.e., go/no-go task). b) An accelerometer attached to the participants hand was used to measure RT, and arm acceleration. The image displays a single participant’s data in the Withdraw condition (one trial, and the corresponding RT and peak acceleration is displayed in red). Zero acceleration indicates that hand velocity was constant. Here, positive acceleration indicates that the arm was moving away from the screen at increasing velocity. RT latency corresponds to the time point where arm acceleration first deviates from zero. Peak positive acceleration refers to the point of time where the participant’s withdrawal velocity is increasing at the fastest rate. Negative acceleration indicates that the participant’s hand movement is slowing down. c) Scatter density plot showing arm peak acceleration as a function of RT (data: whole sample of withdraw condition of the main behavioral experiment, each dot represents a single experimental trial). Photographs in panel a: Common European adder, Vipera berus, by Benny Trapp (https://commons.wikimedia.org/wiki/File:Benny_Trapp_Vipera_berus.jpg), and the corresponding author of the manuscript (based on a photo by Nelli Kraufvelin).

While theories typically assume that threat-relevant stimuli produce responses with shorter reaction times (RTs) than non-threatening stimuli, threat-related stimuli could also trigger more vigorous arm movements (Shadmehr & Ahmed, 2020). From an evolutionarily point-of-view, it would be advantageous to withdraw from threats such as snakes both with low reaction latency and with high acceleration to increase the chances of avoiding a snake bite. We measured behavioral responses using an accelerometer attached to the participant’s hand which allowed us to test, not only whether different animal stimuli evoke different RT, but also test if specific animal classes trigger more vigorous responses than others. If the brain initiates reflexive avoidance of potentially threatening stimuli, participants should withdraw faster (faster RTs and acceleration) from snakes, and possibly also spiders (Davey, 1995), compared to birds and butterflies (which humans rarely report fearing). For the same reason, it might also take longer for participants to approach snakes, or spiders than birds, or butterflies. In a subsequent experiment, we also tested if ERP correlates of attention towards a threat-relevant stimuli (e.g., P1 and EPN) correlate with behavioral responding to threats in the new “naturalistic” paradigm.

## Methods

### Participants

Experiment 1: Thirty-one participants took part in Experiment 1 (7 men, 24 women; age 18–36 years old). Because effect size could not be estimated based on previous studies using power analysis, we estimated sample size based on a sensitivity analysis, and aimed to collect 25 participants, which provides 80% power to observe effects size dz = 0.58 based on a two-tailed comparison of paired means with 0.05 alpha level (Faul et al., 2007). The resulting effect size equals approximately 20 ms difference in RTs assuming SD = *∼*35 ms. Three participants were excluded from the analysis (2 because they were missing either the Withdraw or Approach condition, and one because her long fingernails impeded data collection). Control Experiment: 25 right-handed participants (6 men; age 20–39 years old, and handedness). One participant was excluded from the analysis, because she had no correct answers in the experiment. EEG experiment: Thirty right-handed participants (5 male, age 19–38 years) took part in the experiment. Three participants were excluded from the analysis due to noisy EEG. Given that EPN is a robust correlate of threat detection (dz > 1; e.g. (Grassini et al., 2016, 2019)), the resulting sample size gives sufficient power to detect it. Participants in all studies had no diagnosed neurological disorders, and reported either normal or corrected to normal vision. Participants gave written informed consent before participation in the study, and received study credits for participation. The data was collected at the Department of Psychology and Speech Language Pathology at University of Turku. The study was conducted according to the principles of the Declaration of Helsinki, and was approved by the Ethics Committee for Human Sciences at the University of Turku.

### Stimuli and procedure

The stimuli included photos of snakes, spiders, birds, and butterflies, and photos of mushrooms, pinecones, leaves, and flowers. Each category included 28 different stimuli. The stimuli were collected from various open online sources (e.g., Wikipedia). The luminance histograms of all stimuli (each RGB layer) were matched with the SHINE toolbox (Willenbockel et al., 2010) so that differences in RTs, or ERP amplitudes are not likely to be related to differences in stimulus luminance. Spatial frequency distribution of stimuli was not equated because this manipulation would also likely reduce visibility of the cues the visual system uses to detect, e.g., snakes (Beligiannis et al., 2022). The image resolution was 600 × 450 pixels (37,5 cm × 28 cm on the display, corresponding to approximately 46° × 34° in visual angle). The object displayed on the image was roughly similar in size between different animal conditions. The object was located near the center of the image, and it was clearly visible against the background.

The experiment consisted of a go/no-go task, where participants were seated in front of a large touch screen monitor (Phillips Signage Solutions Multi-Touch Full HD 55” monitor with a 1920 x 1080 resolution, and 60 Hz refresh rate). The experiment consisted of two response conditions: Approach and Withdrawal. In the Approach condition, participants were asked to touch the screen with their finger every time an animal was shown. At the beginning of each trial, the participant kept his/her hand on a platform in front of the touch screen monitor. In the Withdraw condition, participants were asked to hold their finger on the screen (near the center of the screen where the animal may be presented), and withdraw their arm near their torso as soon as they saw an animal. The participants were instructed to respond rapidly, but not to move their hand when no animal was present in the photo. After each trial, the monitor returned to the start screen. The participants were asked to position themselves so that when they extended their arm, they were able to comfortably touch the screen with their finger. Participants performed practice trials of both conditions before the experiment (stimuli used in the practice trials were different that those presented in the main experiment). The Approach and Withdraw conditions each contained 14 different images per stimulus category, presented in random order. Each participant saw each image only once, but stimulus presentation was counterbalanced so that across participants each image was presented in Approach or Withdraw condition with equal probability. The order of the Approach and Withdraw conditions was also counterbalanced across participants. The stimuli were presented using Presentation software (v.22.0/05.10.20). In the Control Experiment, the procedure was identical, but participants responded by pressing (“approach”) or releasing (“withdraw”) a button on a game pad using the thumb of their right hand.

In the main behavioral experiment (but not in the control or EEG experiments), after the participant had completed the behavioral task, he/she was asked to rate on a scale from 0–4 how difficult it would be for them to touch the animal presented in the image: The participant viewed each stimulus (unlimited duration), and reported the rating by pressing a button on the touch-screen monitor. In the instructions, we emphasized, that the participant should consider the specific animal picture that was presented, not the species in general. The highest alternative meant that they would certainly not want to touch the animal, whereas the lowest alternative meant that they could easily touch the animal. We call this variable the “Aversiveness rating”.

At the end of each experiment, we asked participant to rate on a scale from 0 to 4 how much they generally feared snakes, spiders, butterflies and birds (0 = not at all, 4 = very much). Across the experiments, the participants systematically rated higher fear of snakes (M = 1.96, SD = 1.14) and spiders (M = 2.00, SD = 1.12) than birds (M = 0.32, SD = 0.66) and butterflies (M = 0.35, SD = 0.84).

In the EEG Experiment we investigated the association of ERP correlates of threat detection and response times, and so the experiment only included the Withdraw condition (as threat-relevance mainly influenced behavior in this condition). One experimental EEG block included 112 trials (14 stimuli per 8 different stimulus categories). Each participant completed 14–17 blocks in total (i.e., 196–238 trials per condition, per participant). Because the participants performed multiple experimental blocks, in the EEG experiment they were presented with the same stimuli multiple times during the experiment.

### Accelerometer

Arm acceleration was recorded with an MPU6050 accelerometer that was attached on the participant’s wrist with velcro. The sampling frequency of the accelerometer was 100 Hz, and it measured acceleration in three spatial dimensions. The accelerometer is sensitive to the earth’s gravitation, and before each experimental block it was calibrated so that acceleration was 0 m/s^2^ in all three directions. During the calibration, the participant held their hand in a specific position (platform in front of the monitor in the Approach condition, and in the Withdraw condition they held their index finger near the center of the monitor surface) without moving it. The accelerometer was controlled by two Arduino Uno (ATmega328P Arduino Uno R3 AVR® ATmega AVR MCU) units. One of the units was connected to a Dell Latitude E5540 laptop, on which Presentation was run, and the other one was connected to a laptop running a Cool Term win software, which was used to convert the accelerometer output into a text file. Markers indicating stimulus presentation were also printed on the text file with the accelerometer data.

During data analysis, only the dimension of acceleration that measured the approach/withdraw direction was considered (i.e., lateral, and vertical acceleration were not analyzed). The acceleration data was cut to segments corresponding to individual trials, spanning from –200 to 1000 ms relative to stimulus onset. After this, the data was baseline corrected, so that during the prestimulus time-window acceleration was, on average, 0 m/s^2^ for each trial. Baseline correction was necessary because despite the initial calibration in the beginning of each experimental block, small changes in arm angle or rotation changed baseline acceleration values away from 0 m/s^2^.

Peak withdrawal, or approach acceleration was determined for each trial (maximum acceleration in given direction after stimulus onset). Peak withdrawal acceleration was changed to absolute value (to make the negative withdrawal acceleration comparable to positive approach acceleration). RT was determined by searching for the first sample where acceleration began to continuously approach the peak acceleration.

### EEG

EEG was recorded with 32 passive electrodes placed according to the 10-10 electrode system (EasyCap GmbH, Herrsching, Germany). Surface electromyograms (EMGs) were measured with two electrodes placed below and to the side of the left eye. Reference electrode was placed on the nose, and ground electrode on the forehead. EEG was recorded with a NeurOne Tesla amplifier using 1.4.1.64 software (Mega Electronics Ltd., Kuopio, Finland). Sampling rate was 500 Hz.

EEG data was processed using EEGlab software (Delorme & Makeig, 2004) running on Matlab R2020b. First, a marker representing the latency of arm movement onset was added to the data. Here, the movement onset latency was retrieved from the touch screen data, as it measured RT with higher resolution than the accelerometer. This marker also contained the peak acceleration of the arm movement (measured using the accelerometer). After this, EEG data was high-pass filtered (1 Hz), and then low-pass filtered (40 Hz). Bad electrodes were removed, reference was changed to average, the data was epoched, and independent component analysis was run. IC label (Pion-Tonachini et al., 2019) was used to keep independent components with at least 70% probability that the component was brain-based. Outlier trials were rejected using joint probability functions with the criteria of three standard deviations. Finally, removed channels were interpolated back into the data.

Based on previous research (Olofsson et al., 2008b; Smith et al., 2003; Sperl et al., 2021), we analyzed ERPs spanning time-windows from P1 to EPN (this time-window also included the N1 wave). Based on the visual inspection of differences between threat-relevant (mean of snake and spider stimuli), and non-threat conditions (mean of butterfly and bird stimuli), P1 (70–110 ms) and EPN (200–240 ms) analysis was focused on electrodes over the occipital pole (O1, O2, and Iz), and N1 (140–160 ms) analysis was based on lateral occipitoparietal electrodes (P7 and P8) because threat-related modulation was strongest around these electrodes and time-windows (see, Fig. 4b). In addition to analyzing the “peak” P1 amplitude, we analyzed the onset amplitude of P1 to determine if the very earliest visual stimulus evoked amplitudes correlate with threat-detection (50–60 ms; electrodes O1, O2, and Iz). The analysis of this time-window was performed post-hoc, after a reviewer guided the authors towards studies demonstrating that treat-relevance may modulate ERPs already in this time-window corresponding to previously reported effect on the C1 wave (Hintze et al., 2014; Sperl et al., 2021; Stolarova et al., 2006; Thigpen et al., 2017). Average ERP amplitude in the given time window and electrodes was used for data analysis in single-trial manner (see Statistical analysis paragraph). Later time-windows (e.g., LPP) were not examined because activity in these later time-windows overlapped with behavioral responding, making it difficult to tease apart the correlates of threat processing from motor behavior.

### Statistical analysis

RT and peak acceleration analyses were based on trials where the participants made a behavioral response to the presented animal stimulus. The data was analyzed on single-trial level using linear mixed-effects models. For the main behavioral experiment and the control experiment (with button-press/release responding) the model was as follows: *Dependent_variable ∼ Response_condition * Animal_category + RT_or_acceleration + (1 + Response_condition | Participant) + (1 + Response_condition | Stimulus)*, where *Dependent_variable* is either RT or peak acceleration, *RT_or_*acceleration is the z scored RT or acceleration (depending on which was the dependent variable; this term was removed from the statistical analysis of the control experiment), *Response_condition* refers to the Approach vs. Withdraw condition (button press vs. release in the control experiment), *Animal_category* is the species present in the stimulus, and *Stimulus* refers to the specific animal stimulus presented in the given trial. The random-effect structure thus included between-participant variation in response speed (i.e., intercept) as well as in how response condition influenced the behavioral response. In addition, the random-effects structure took into account variation between individual stimuli. This was considered important because differences between photos could potentially influence RT or arm acceleration (e.g., due to differences in the size and orientation of the animal, there may be systematic variation in responses to specific stimuli). This fixed– and random-effect structure led to best models based on Akaike and Bayesian Information Criteria. In the models, the intercept represents the Approach/Butterfly condition. In the EEG experiment, data was analyzed using following linear-mixed effects models: *Dependent_variable ∼ Animal_category + RT + Acceleration + (1|Participant)*, where the dependent variable is the mean P1, N1, or EPN amplitude in the specified time-windows.

Reported aversiveness of the stimulus (which was collected as part of the main behavioral experiment), or participants’ general fear of the stimulus species was not included in the regression models because these predictors were strongly collinear with the *Animal_category* predictor. Instead, we examined if between-participants variation in reported aversiveness predicted RTs and peak arm acceleration (in the main experiment). For each participant, we calculated mean RT, mean peak arm acceleration, and mean reported aversiveness of the presented stimuli for each animal category, and response condition. We then used *Dependent_variable ∼ Response_condition*Aversiveness_rating + Animal + (1 + Response_condition |Participant)* regression models to investigate if aversiveness of the animal predicted behavioral responses (interactions between specific animal species and aversiveness did not approach significance so they were left out of the model). In the EEG experiment, following a similar procedure we examined if between-participant variation in mean RT, mean peak acceleration, or the participant’s fear rating predicted ERP amplitude (*Dependent_variable ∼ Animal + RT + peak_acceleration + (1|Participant)*).

The data and the analysis scripts are available at https://osf.io/j6sqx/.

## Results

### Main Behavioral Experiment

To characterize the participants’ behavioral responses to the stimuli, we calculated RT latency and peak arm acceleration for each trial (Fig. 1b). We used linear mixed-effects models to test if trial-by-trial variation in the behavioral responses were modulated by the stimulus presented (i.e., animal species pictured), or how the participant responded (withdraw vs. approach movement). The results are visualized in Fig. 2. Participants’ responses were highly accurate (Fig. 2, left column), and displayed only minor, and not statistically significant differences between different the experimental conditions. The results of a model predicting variation in RTs are shown in Table 1 (upper part). The intercept term shows that across participants, mean RT to butterfly stimuli in the Approach Condition was 304 ms. Lack of main effect of different animal categories indicates that in the Approach condition, participants’ RTs were not statistically significantly modulated by the species they responded to. In the Withdraw condition, RTs were on average 95 ms slower than in the Approach condition. Crucially, interactions between Withdraw condition and Snake/Spider animals shows that when participants withdrew from the image, these two types of images were associated with about 19 ms (t = –2.60) and 27 ms (t = –3.77) speed-up of RTs, respectively. This result shows that participants *withdrew* from snakes and spiders with faster RT latency (Fig. 2, middle column) as compared to butterflies and birds. Peak acceleration was negatively associated with RT latency: Increase in peak arm acceleration corresponding to standard deviation was associated with about 5.5 ms faster RTs.

**Figure 2.**
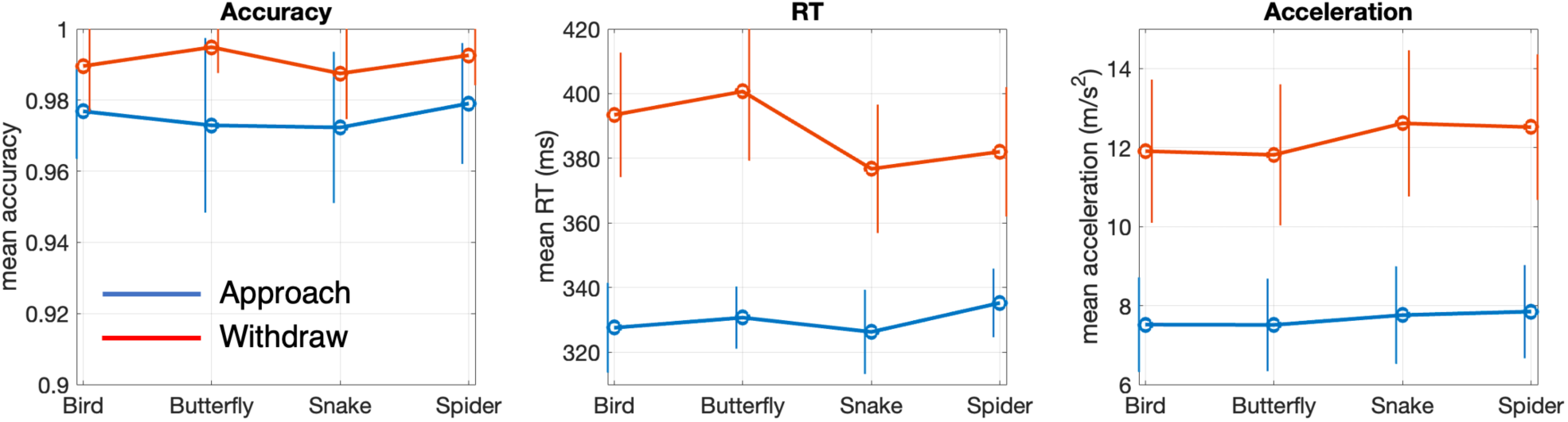
Behavioral results. The Approach condition is visualized in blue, and the Withdraw condition in red. Columns show response accuracy, RT, and peak acceleration, respectively. Error bars display the 95% CI.

**Table 1.**
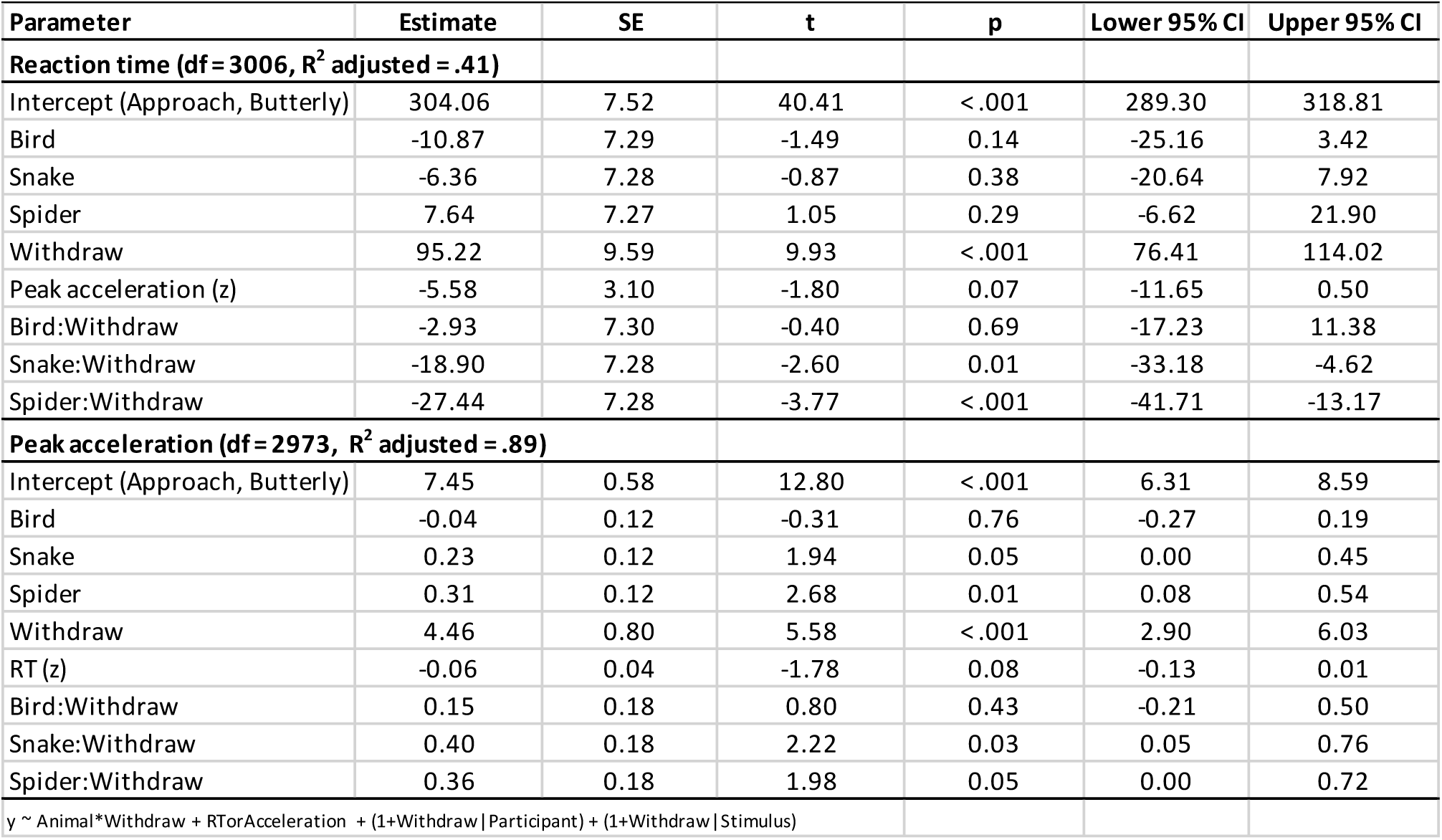
Results of mixed-effects regression analysis (Main behavioral experiment)

|  |  |  |  |  |  |  |
| --- | --- | --- | --- | --- | --- | --- |
| Parameter | Estimate | SE | t | p | Lower 95% CI | Upper 95% CI |
| **Reaction time (df = 3006, R<sup>2</sup> adjusted = .41)** |  |  |  |  |  |  |
| Intercept (Approach, Butterfly) | 304.06 | 7.52 | 40.41 | < .001 | 289.30 | 318.81 |
| Bird | -10.87 | 7.29 | -1.49 | 0.14 | -25.16 | 3.42 |
| Snake | -6.36 | 7.28 | -0.87 | 0.38 | -20.64 | 7.92 |
| Spider | 7.64 | 7.27 | 1.05 | 0.29 | -6.62 | 21.90 |
| Withdraw | 95.22 | 9.59 | 9.93 | < .001 | 76.41 | 114.02 |
| Peak acceleration (z) | -5.58 | 3.10 | -1.80 | 0.07 | -11.65 | 0.50 |
| Bird:Withdraw | -2.93 | 7.30 | -0.40 | 0.69 | -17.23 | 11.38 |
| Snake:Withdraw | -18.90 | 7.28 | -2.60 | 0.01 | -33.18 | -4.62 |
| Spider:Withdraw | -27.44 | 7.28 | -3.77 | < .001 | -41.71 | -13.17 |
| **Peak acceleration (df = 2973, R<sup>2</sup> adjusted = .89)** |  |  |  |  |  |  |
| Intercept (Approach, Butterfly) | 7.45 | 0.58 | 12.80 | < .001 | 6.31 | 8.59 |
| Bird | -0.04 | 0.12 | -0.31 | 0.76 | -0.27 | 0.19 |
| Snake | 0.23 | 0.12 | 1.94 | 0.05 | 0.00 | 0.45 |
| Spider | 0.31 | 0.12 | 2.68 | 0.01 | 0.08 | 0.54 |
| Withdraw | 4.46 | 0.80 | 5.58 | < .001 | 2.90 | 6.03 |
| RT (z) | -0.06 | 0.04 | -1.78 | 0.08 | -0.13 | 0.01 |
| Bird:Withdraw | 0.15 | 0.18 | 0.80 | 0.43 | -0.21 | 0.50 |
| Snake:Withdraw | 0.40 | 0.18 | 2.22 | 0.03 | 0.05 | 0.76 |
| Spider:Withdraw | 0.36 | 0.18 | 1.98 | 0.05 | 0.00 | 0.72 |
| y ~ Animal*Withdraw + RTorAcceleration + (1+Withdraw\|Participant) + (1+Withdraw\|Stimulus) |  |  |  |  |  |  |

Next, we examined if the target animal species, or the type of behavioral response predicted peak arm acceleration (Table 1, lower part). The intercept represents the Approach/Butterfly condition, indicating that peak response acceleration was on average 7.4 m/s^2^. The main effects of snake and spider animal categories indicate that participants’ approach responses to these species evoked responses with 0.23 m/s^2^ (t = 1.94), and 0.31 m/s^2^ (t = 2.68) higher peak acceleration, respectively. Interaction effects between these two species and Withdraw condition indicate that in the Withdrawal condition, arm acceleration was further increased by 0.40 m/s^2^ (t = 2.22), and 0.36 m/s^2^ (t = 1.98) for snakes and spiders respectively. This means that altogether based on the model, in the Withdraw condition arm acceleration was 0.63 m/s^2^, and 0.67 m/s^2^ faster for snakes and spiders when compared to butterflies. In general, arm acceleration was 4.4 m/s^2^ slower in the Approach condition (likely because participants had to restrict the velocity of their arm in order to not hit the touch screen monitor), and one SD decrease in RT predicted 0.06 m/s^2^ faster arm acceleration.

So far, the results show that within participants, snakes and spiders evoke faster withdrawal RTs and higher arm acceleration than birds of butterflies. Next, we examined if subjectively experienced aversiveness of the animal species predicts variation *between participants* (in RT, and peak hand acceleration). First, as expected, the participants reported low aversiveness for the bird and butterfly stimuli, but clearly higher aversiveness for snakes and spiders (Supplementary Figure 1). Second, the results showed that while aversiveness of the animal did not predict approach RTs (t = –0.64) or acceleration (t = 0.021), in the withdraw condition, higher aversiveness was associated with faster RTs (8.8 ms faster responses per unit increase in rating, t = –4.00), and higher arm acceleration (0.14 m/s^2^ higher acceleration per unit increase in rating, t = 2.34). The results are visualized in Supplementary Figure 1, and the full results of the regression model are presented in Supplementary Table 1.

### Control Experiment

To determine if the naturalistic paradigm was curial for observing the behavioral results, we next tested if similar results are obtained when RT is measured using a button press/release task. In this control experiment, participants were instructed to respond to the stimulus either by pressing a button (mimicking the Approach condition), or by releasing the button (mimicking the Withdrawal condition). The results (Supplementary Fig. 2) showed that while accuracy was very high in general, the participants made somewhat more mistakes in snake and spider conditions (compared to bird and butterfly conditions; however, this effect was driven by three outlier participants). As shown in Supplementary Table 2, RTs revealed no statistically significant differences between different animal conditions in either button press, or release conditions, and in fact, participants responded fastest to bird stimuli (in both button-press and release conditions). This indicates that a naturalistic research setting was necessary for detecting rapid behavioral responses to threat-relevant stimuli.

### EEG Experiment

One major shortcoming of previous research on the ERP correlates of threat detection is that the studies have not tested whether ERP correlates of attention towards threat-related stimuli predict behavioral responding such as RTs. We tested the electrophysiological correlates of naturalistic withdrawal responses by combining the behavioral paradigm with EEG (only the Withdraw condition). In general, the behavioral results (Fig. 3) replicate the earlier findings: Withdrawal RTs were fastest to snakes, although the size of the effect (12.3 ms) was about half of what was observed previously. Unlike in the previous experiment, participants did not withdraw more rapidly from spiders. Peak arm withdrawal acceleration was higher for both snakes and spiders (than butterflies, or birds), and the size of the effect is comparable to the main behavioral experiment. The full behavioral results are displayed in Supplementary Table 3.

**Figure 3.**
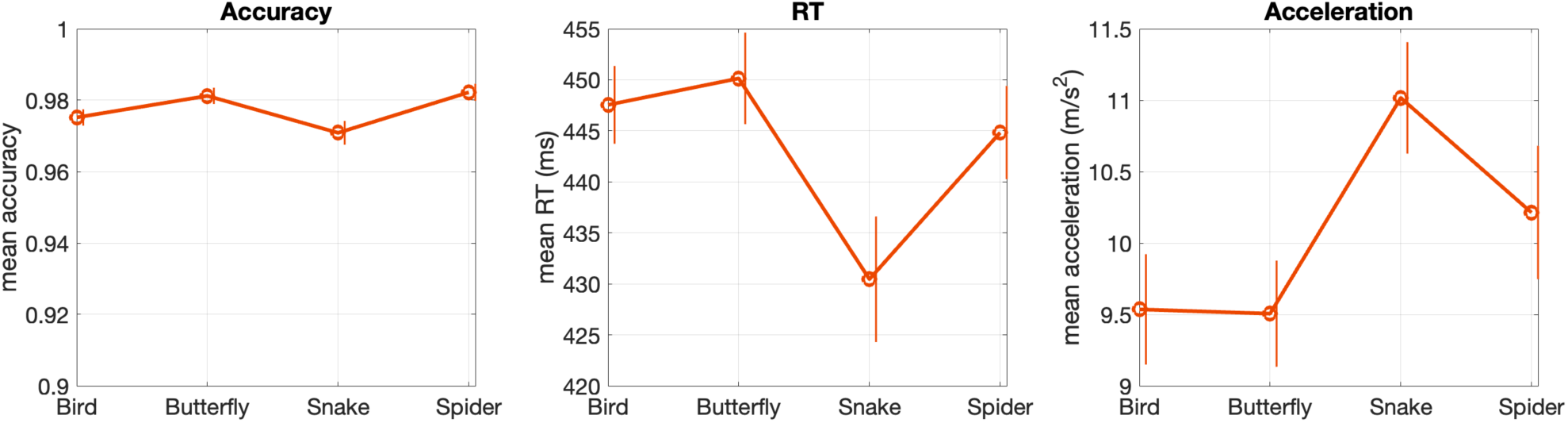
Behavioral results of the EEG experiment. Error bars display 95% CI.

Similar to the main behavioral results, between-participant variation fear ratings predicted withdrawal RTs and arm peak acceleration: Higher reported fear was associated with faster RTs (β = –1.41 per unit increase in fear rating, t = –2.11), and faster arm acceleration (β = 0.13 per unit increase in fear rating, t = 2.60).

As shown in Fig. 4a, the animal images evoked ERPs with an onset around 50 ms, after which prominent P1 (80–100 ms), and N1 (120–140 ms) waves in the occipital electrodes were observed. These were followed by a series of waves between 200–500 ms. To understand how the threat-related stimuli modulated ERPs, we first visualized general differences in ERPs to threat-relevant vs non-threat-relevant images (Fig. 4b). This revealed three time-windows where threat-relevance seemed to modify ERP amplitudes: Threat-related stimuli were associated with changes in P1 (peak around 90 ms after stimulus onset) and N1 (peak around 150 after stimulus onset) amplitudes, and threat-related images produced more negative amplitudes between 200–300 ms, corresponding to the EPN. As shown in Fig. 4c, arm withdrawal RT strongly influenced ERPs between 240–800 ms, and as previously reported (Delorme et al., 2007; Makeig et al., 2004), movement produced a clear time-varying positive wave whose timing corresponded to the RT.

**Figure 4.**
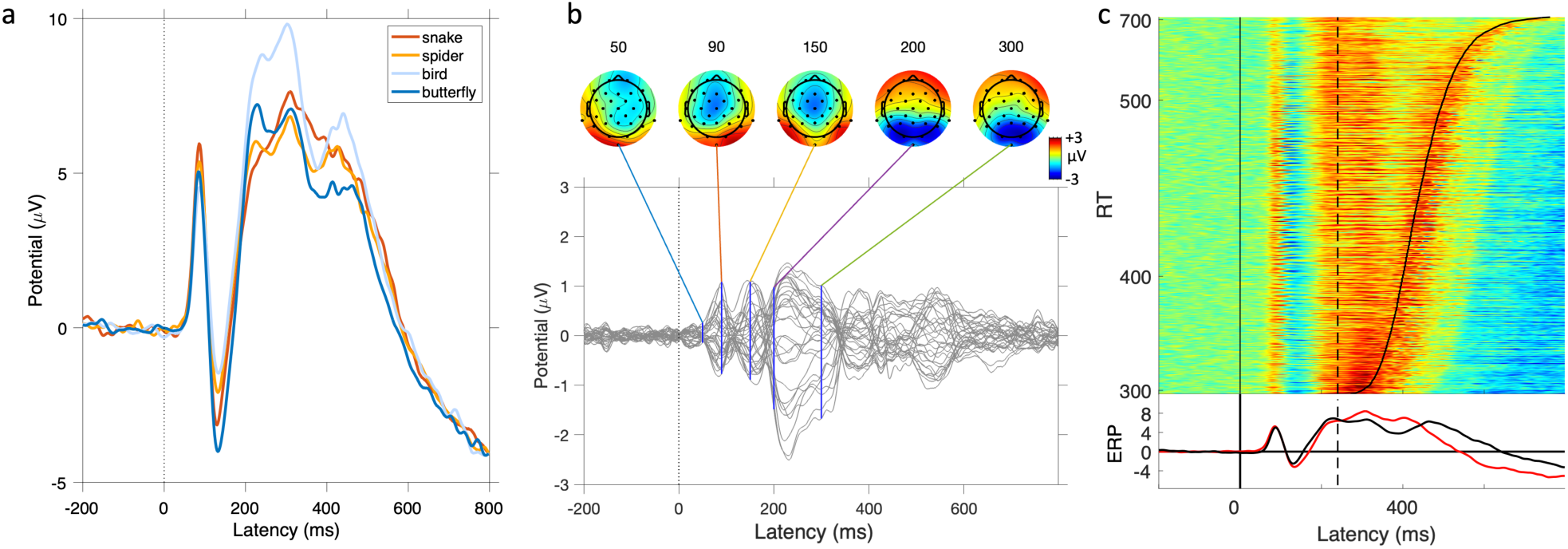
Grand average ERPs. a) ERPs to different animal categories (mean of channels O1, O2, and Iz). b) Difference wave between threat-relevant (snake and spider) vs. non-treat-relevant images (bird, and butterfly), and the corresponding scalp maps (of the difference wave) at specified latencies. Each line represents one channel. c) ERP-image visualizing how arm movement influenced single-trial ERPs (mean of channels O1, O2, and Iz). Each experimental trial has been color coded (red = positive amplitude, blue = negative), and the trials have been sorted according to RT. Vertical solid line at 0 ms indicates the onset of the visual stimulus. The arched solid line shows the RT in the corresponding trials. Dashed vertical line shows the 240 ms time point corresponding to the end of the EPN time-window. The ERP-image has been smoothed by Gaussian moving average (SD = 5 trials). Grand average ERP is presented below the ERP-image. The red line represents the ERP of the fastest 50% of trials, and the black line the slowest 50% of trials.

We used linear mixed-effects models to test if animal species and the (z scored) behavioral metrics (RT and peak acceleration) predicted P1 onset amplitudes (channels O1, O2, and Iz; time-window: 50–60 ms), P1 “peak” amplitude (channels O1, O2, and Iz; time-window: 70–110 ms), N1 amplitude (channels P7 and P8; time-window: 120–140 ms; please see Supplementary Figure 3 for ERPs in electrodes P7 and P8), and EPN amplitude (channels O1, O2, and Iz; time-window: 200– 240 ms). Because the models did not reveal interactions between animal category and RT, or peak acceleration, the interaction terms were removed from the final models.

The results are presented in Table 2. Figure 5 shows the association between ERP amplitudes, and RT (upper panels) and acceleration (lower panels). Birds evoked the weakest activation in the P1 onset time-window (t = –4.28), but amplitudes evoked by snake or spider images did not differ statistically significantly from the butterfly images. Stronger P1 onset amplitude predicted faster RTs (t = –5.80), but slower arm withdrawal acceleration (t = –2.15).

**Figure 5.**
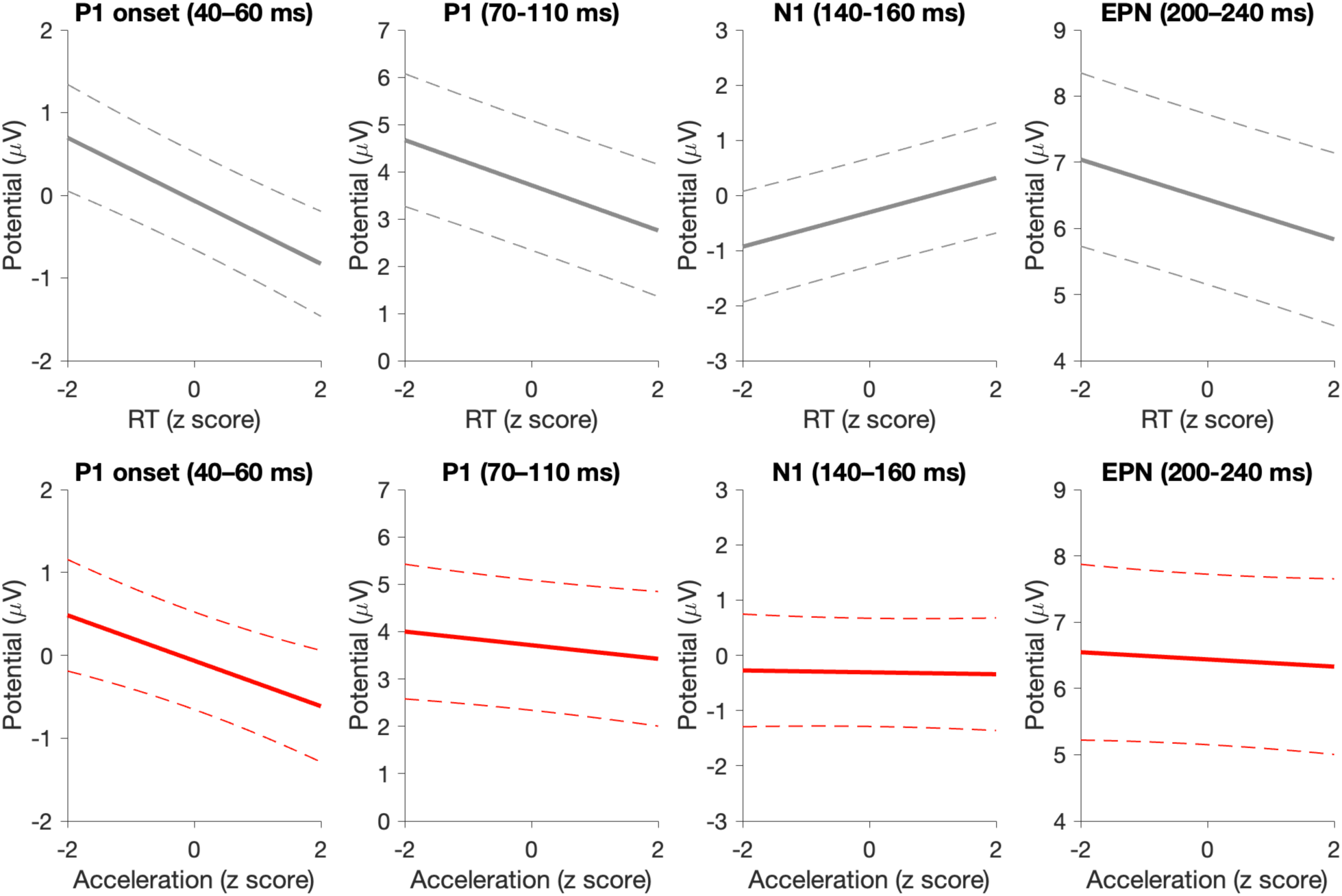
Correlation between RT (upper panels) and acceleration (red lines, lower panels) with C1, P1, N1, and EPN amplitudes (results of the regression model presented in Table 5). The dashed line is the 95% CI.

**Table 2.**
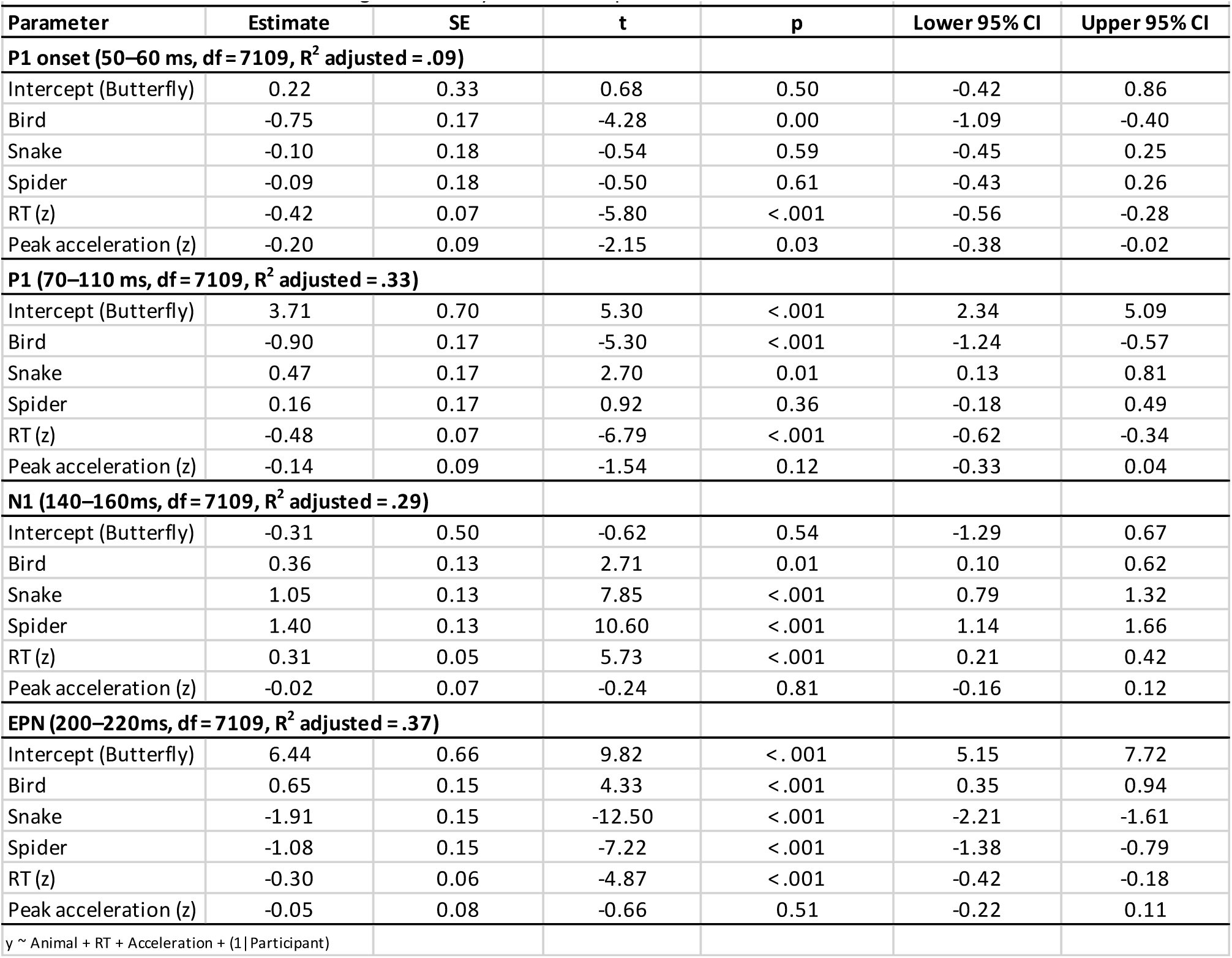
Results of the mixed-effects regression analysis on ERP amplitudes.

|  |  |  |  |  |  |  |
| --- | --- | --- | --- | --- | --- | --- |
| Parameter | Estimate | SE | t | p | Lower 95% CI | Upper 95% CI |
| **P1 onset (50–60 ms, df = 7109, R<sup>2</sup> adjusted = .09)** |  |  |  |  |  |  |
| Intercept (Butterfly) | 0.22 | 0.33 | 0.68 | 0.50 | -0.42 | 0.86 |
| Bird | -0.75 | 0.17 | -4.28 | 0.00 | -1.09 | -0.40 |
| Snake | -0.10 | 0.18 | -0.54 | 0.59 | -0.45 | 0.25 |
| Spider | -0.09 | 0.18 | -0.50 | 0.61 | -0.43 | 0.26 |
| RT (z) | -0.42 | 0.07 | -5.80 | <.001 | -0.56 | -0.28 |
| Peak acceleration (z) | -0.20 | 0.09 | -2.15 | 0.03 | -0.38 | -0.02 |
| **P1 (70–110 ms, df = 7109, R<sup>2</sup> adjusted = .33)** |  |  |  |  |  |  |
| Intercept (Butterfly) | 3.71 | 0.70 | 5.30 | <.001 | 2.34 | 5.09 |
| Bird | -0.90 | 0.17 | -5.30 | <.001 | -1.24 | -0.57 |
| Snake | 0.47 | 0.17 | 2.70 | 0.01 | 0.13 | 0.81 |
| Spider | 0.16 | 0.17 | 0.92 | 0.36 | -0.18 | 0.49 |
| RT (z) | -0.48 | 0.07 | -6.79 | <.001 | -0.62 | -0.34 |
| Peak acceleration (z) | -0.14 | 0.09 | -1.54 | 0.12 | -0.33 | 0.04 |
| **N1 (140–160ms, df = 7109, R<sup>2</sup> adjusted = .29)** |  |  |  |  |  |  |
| Intercept (Butterfly) | -0.31 | 0.50 | -0.62 | 0.54 | -1.29 | 0.67 |
| Bird | 0.36 | 0.13 | 2.71 | 0.01 | 0.10 | 0.62 |
| Snake | 1.05 | 0.13 | 7.85 | <.001 | 0.79 | 1.32 |
| Spider | 1.40 | 0.13 | 10.60 | <.001 | 1.14 | 1.66 |
| RT (z) | 0.31 | 0.05 | 5.73 | <.001 | 0.21 | 0.42 |
| Peak acceleration (z) | -0.02 | 0.07 | -0.24 | 0.81 | -0.16 | 0.12 |
| **EPN (200–220ms, df = 7109, R<sup>2</sup> adjusted = .37)** |  |  |  |  |  |  |
| Intercept (Butterfly) | 6.44 | 0.66 | 9.82 | <.001 | 5.15 | 7.72 |
| Bird | 0.65 | 0.15 | 4.33 | <.001 | 0.35 | 0.94 |
| Snake | -1.91 | 0.15 | -12.50 | <.001 | -2.21 | -1.61 |
| Spider | -1.08 | 0.15 | -7.22 | <.001 | -1.38 | -0.79 |
| RT (z) | -0.30 | 0.06 | -4.87 | <.001 | -0.42 | -0.18 |
| Peak acceleration (z) | -0.05 | 0.08 | -0.66 | 0.51 | -0.22 | 0.11 |
| y ~ Animal + RT + Acceleration + (1\|Participant) |  |  |  |  |  |  |

Snakes produced the strongest P1 waves (t = 2.7, compared to butterflies), and birds were associated with the smallest P1 amplitudes (t = –5.3, compared to butterflies). Faster RTs were associated with stronger P1 amplitude. Peak acceleration was not statistically significantly associated P1 amplitude.

In the N1 time-window, both snakes (t = 7.8, compared to butterflies) and spiders (t = 10.6, compared to butterflies) were associated with decreased (i.e., more positive) N1 amplitudes. Yet, across stimuli, faster RTs were associated with stronger (i.e., more negative) N1 amplitudes (Fig. 5, middle panel). Peak acceleration was not statistically significantly associated N1 amplitude.

Finally, as expected, in the EPN time-window, snakes and spiders produced stronger EPN than butterflies, and birds evoked the least strong EPN (t values ≤ 4.3). However, interestingly, EPN amplitude was inversely correlated with RT: The stronger the EPN (i.e., the more negative the amplitudes), the slower the RT (Fig. 5, right panel). This is at odds with the common assumption that EPN reflects processes that mediate rapid behavioral avoidance by allocating attention towards threatening animals.

It could be argued that, although the EPN time-window (200–240 ms) preceded the motor responses (Fig. 4c), preparatory motor activation preceding the fastest RTs nevertheless conflate the measured EPN. To rule out this possibility, we excluded from the analyses RTs that were shorter than 420 ms. This way, any behavioral response took place a minimum of 200 ms after the EPN time-window. Despite this, P1 onset (t = –2.64), P1 “peak” (t = –4.1), N1 (t = 2.7), and EPN (t = – 2.3) amplitudes showed the same modulation by RT. This shows that the negative correlation between EPN and RT was not due to a motor confound. In the control analysis peak acceleration was not statistically significantly associated with EPN amplitude in any of the time-windows.

We also investigated if, across participants, mean ERP amplitudes (P1 onset, P1, N1, and EPN) correlated with participants’ mean withdrawal RT and peak acceleration. The results (Supplementary Figure 4) did not reveal any significant correlations. This means that while P1, N1, and EPN correlate with RT and arm withdrawal acceleration within participants, similar association is not observed between participants. Finally, higher fear rating predicted stronger N1 amplitudes (Estimate = –0.24, t = –2.76), but fear was not associated statistically significantly with P1 onset (t = 0.40), P1 (t = –0.91), or EPN (t = –1.18) amplitudes.

## Discussion

We have shown that animals humans often report fearing (snakes and spiders) can trigger rapid behavioral withdrawal responses when compared to non-threatening species (butterflies and birds). In addition to shorter withdrawal RTs, the threat-relevant animals triggered faster arm acceleration. This speed-up of behavioral responding was only observed in a “naturalistic” paradigm, where individuals responded using arm movements. This suggests that the neural mechanisms that enable rapid withdrawal from threats do not constitute a “modality general” speed-up of all behavioral responding, but are specific to certain types of behavioral responses that need to be measured using a naturalistic paradigm. Across participants, the speed of the withdrawal correlated with how aversive the participants experienced the animal species used as stimuli, suggesting that the speed of the withdrawal reaction also reflects trait-level differences between individuals. This suggests that the withdrawal speed could provide information about the possible relationship between pathologically active or inactive threat detection circuits, and neuropsychiatric condition (such as anxiety disorders, or psychopathy; (Lake et al., 2011; McFadyen et al., 2020; Taschereau-Dumouchel et al., 2022)). In contrast, with the exception of N1 amplitude, the ERP correlates of threat detection did not covary statistically significantly with participants’ fear of the target stimulus across participants, and between participant variation in ERP amplitudes did not predict RT or acceleration differences across participants.

Because our results show that rapid behavioral responding is linked specifically with withdraw movements, the results support the interpretation that the responses were mediated by circuits that mediate escape from threats. In addition to being based on “naturalistic movements”, we suggest that a key factor in the present paradigm is that it mimicked *interaction* with the stimuli. If our interpretation holds, then the speed-up of behavioral withdrawal may only be observed when the participant is close enough to the stimulus, as was the case in the present study. Studies on non-human animals using naturalistic experimental paradigms suggest that basolateral amygdala activation strongly correlates with the initiated behavioral response (e.g., its velocity), but also with the proximity of the threat (Amir et al., 2015; Choi & Kim, 2010). We assume that by incorporating more “naturalistic” threat elements to the present paradigm (e.g., moving, or looming stimuli (Yilmaz & Meister, 2013)), the paradigm could reveal even stronger evidence for automatic withdrawal from threats. Our results support the findings reported by previous naturalistic studies on threat detection in humans (Rinck et al., 2010, 2021), but also extend these by showing that the speed-up of behavioral responding to threats may be movement specific.

We utilized an accelerometer to measure behavioral responses to the stimuli, which allowed us to examine not only RTs, but also if arm acceleration was modulated by the threat-relevant stimuli. From an evolutionary perspective, it would make sense that animals withdraw from threatening stimuli, not just with rapid latency, but also with high acceleration, to maximize the chances of escape. Our results show that although RT and arm peak acceleration correlated, the effects of threat on RT and arm peak acceleration could be dissociated: Whereas the effect of threat on RT was solely linked to withdrawal responses, hand acceleration was modulated by threat also during approach behavior (although also arm acceleration to threats was further increased when participants withdraw from the stimuli).

Within participants, the latency of the behavioral withdrawal reaction was associated with electrophysiological signatures that are typically interpreted as correlates of selective attention. Fast RTs were also associated with stronger early brain responses in the P1 time-window (70–140 ms after stimulus onset), including the very onset of P1 activation (40–60 ms), although threat-relevance of the stimuli did not modulate amplitudes in the P1 onset time window. Unexpectedly, stronger P1 onset amplitudes were associated with slower arm withdrawal acceleration.

Similar to P1, stronger N1 was associated with faster RTs in the present study, even though threat-related stimuli evoked decreased N1 amplitudes (compared to neutral stimuli) in the present study, suggesting that processes that allocate attentional priority to threat-related stimuli, and mediate behavioral responding, can be dissociated. Given that scalp recorded ERPs reflect the summed activation of number of brain sources, we suggest that the decreased N1 amplitudes for threat-relevant stimuli may reflect the same process that amplifies the P1 potential (i.e., the P1 “threat-effect” may continue longer than the P1 wave visible on scalp recorded ERPs). The contributions of different underlying ERP sources could be disentangled, for example, using source separation approaches such as independent component analysis (Colombari & Railo, 2023; Onton et al., 2006).

The main source of the P1 localizes to early visual cortices (Carretié et al., 2022; di Russo et al., 2002), suggesting that that relatively low-level visual processes signaled the presence of a potentially threatening stimulus when a snake was presented. In line with this, while rapid unconscious responses to threat-relevant stimuli are sometimes attributed to subcortical structures(Evans et al., 2018; Salay et al., 2018; Wei et al., 2015), recent studies indicate that rapid cortical processing also plays a key role in threat processing (Carretié et al., 2022; Li & Keil, 2023; Miskovic & Keil, 2012; West et al., 2011). Given that the present stimuli were not designed to measure the C1 wave, it remains open whether the initial wave of visual stimulus driven cortical activity is sensitive to threat-relevance and predicts behavioral responding, as suggested by fear conditioning studies (Hintze et al., 2014; Sperl et al., 2021; Stolarova et al., 2006; Thigpen et al., 2017). The P1 typically precedes the earliest correlate of conscious visual perception (Förster et al., 2020), suggesting that snake detection may be mediated by unconscious perception. However, since the animal stimuli in the present study were consciously perceived, the present results do not demonstrate that unconscious processes alone are sufficient to explain fast behavioral withdrawal from snake images. Research suggests that both fast RTs (Railo et al., 2015) as well as responding to snake stimuli may require conscious perception (Cox et al., 2018; Grassini et al., 2016).

In the present EEG experiment, snake stimuli elicited the fastest behavioral responses and showed the earliest amplified ERP correlates. In the main behavioral experiment, both spiders and snakes evoked comparably rapid withdrawal responses. This difference in the results of the two experiments may be related to methodological differences between the experiment. The EEG experiment was considerably longer, only included the Withdraw condition (also, RTs were generally slower in the EEG experiment), and participants viewed the same stimuli multiple times during the EEG experiment. Previous studies using schematic stimuli have shown that also spider stimuli may evoke enhanced responses during the earliest stages of cortical processing (Carretié et al., 2022).

A key novel finding in the present study is that the amplitude of the P1 response positively correlated, not just with snake “detection”, but also with behavioral withdrawal RT. This suggests that the same mechanism that gives threat-relevant stimuli attentional priority early on in visual processing also influences the speed of behavioral withdrawal. Similar to P1, higher N1 amplitudes were associated with faster RTs, in line with the interpretation that “stronger” stimulus-evoked activity was associated with faster RTs. The correlation between ERPs and withdrawal RTs was observed across all animal stimuli, suggesting that the neural mechanisms that enable fast withdrawal responses are not specific to snakes. Rather, snake stimuli engaged this early amplification mechanism that increased the probability of fast behavioral withdrawal. Consistent with this, P1 amplitudes correlate with multiple different types of threats (Brown et al., 2010).

Contrary to prevailing theories (and contrary to the effects observed in P1 and N1 time windows), stronger EPN amplitude (200–240 ms after stimulus onset) was associated with slower withdrawal RTs. Given the similarities to an ERP correlate of feature-based attention known as selection negativity (Hillyard & Anllo-Vento, 1998; Keil & Müller, 2010), EPN may be interpreted as a sign of attention towards the features of the threat-relevant stimulus. EPN could be described as a correlate of top-down attention to the threat, whereas P1, for example, reflects a more stimulus-driven “bottom-up” capture of attention. The negative correlation between EPN amplitude and RT suggests that rapid withdrawal from threat-related stimuli engages multiple attentional components that differentially contribute to withdrawal speed, and that selective attention to the threat-relevant stimulus may not always be beneficial for response speed.

Conscious perception of visual stimuli correlates with activity in a similar time-window and electrodes as EPN (Förster et al., 2020), and Grassini et al. have shown that snake and spider images only elicit EPN when participants consciously perceive the stimuli (Grassini et al., 2016). This means that EPN may reflect processes related to conscious perception of stimulus details. This suggests that rapid withdrawal responses are based on automatic, reflex-like processing, but attention to consciously discriminate the details of the stimulus may interrupt this automatic process, and consequently delay fast withdrawal. Our results concerning the association between ERP amplitudes and behavioral responding are correlative, but because the ERP correlates preceded the behavioral responses, the ERPs may also be causally linked to the speed-up of behavioral responding.

Our results therefore suggest that the link between attention towards the threat-relevant stimulus and behavioral responding is not straightforward. If our interpretation is correct, rapid withdrawal from a threat requires not only early prioritization of the threat-relevant stimulus, but also rapid disengagement of attention from the threat. Also context likely modulates the likelihood that a person keeps engaged with the threat-relevant stimulus. For example, Rinck et al. (2010) showed that when individuals who fear spiders encountered a spider when walking through a virtual reality museum, they gazed at the spider longer than participants who did not report fearing spiders (i.e., they kept engaged with the threat-relevant stimulus). In this context, the individual may prefer to keep the threat-relevant spider in the field-of-view to make sure he can maintain distance to it. Research suggests that heightened anxiety is associated with difficulty in disengaging from the stimulus (Fox et al., 2001; Yiend & Mathews, 2001). Our results, however, indicate that the higher the individual’s aversiveness of the target animal, the faster they also withdrew from it. We attribute this finding to the fact that the picture of the threat-related animal was near the participant’s hand in the present study, meaning that participants who feared e.g., spiders, likely also wanted to distance themselves from them.

### Conclusions

Our results show that images of snakes and spiders elicit fast and vigorous arm withdrawal in humans. Because similar effect was not observed with approach movements, or button-press/release responding, the results suggest snake and spider stimuli activated neural circuits associated specifically with escape and avoidance. Our electrophysiological results suggest that the fast withdrawal was based on two opposite attentional processes that preceded the behavioral response. First, presence of threat was signaled by early amplification of the stimulus-related activation, which also correlated with faster behavioral withdrawal. Second, while threat-related stimuli also activate further top-down attention to the details of the stimulus, failure to rapidly “disengage” attention from the threat was associated with slower withdrawal reactions. This suggests that rapid behavioral withdrawal is mediated by automatic reflex-like circuits which may be interrupted by attentive processing. We suggest that the present study describes a fruitful approach to disentangle the processes that underlie behavioral responses to threats. For example, the present approach could be easily adopted to study responses in children, or even in non-human primates, and the paradigm could be combined with experimental manipulations such as attentional cuing (e.g., Fox et al., 2001; Yiend & Mathews, 2001), fear conditioning (e.g., Hintze et al., 2014; Sperl et al., 2021; Stolarova et al., 2006; Thigpen et al., 2017), or manipulations of stimulus properties (e.g., Méndez-Bértolo et al., 2016; Vuilleumier et al., 2003) to better understand the neural and cognitive underpinnings of behavioral responses towards threat-relevant stimuli.

## Acknowledgements

We thank Clara-Theresia Kolehmainen for help with data collection.

## Author contributions

H.R.: designed the study, analyzed the data, wrote the manuscript, and supervised data collection.

T.L. wrote the experimental scripts and build the apparatus to measure behavioral responses with accelerometer. N.K. Collected data, and analyzed the EEG data. J.S. Collected data.

## SUPPLEMENTARY INFORMATION

*Railo et al. — Rapid withdrawal from a threatening animal is movement-specific and mediated by reflex-like neural processing*

**Supplementary Figure 1.**
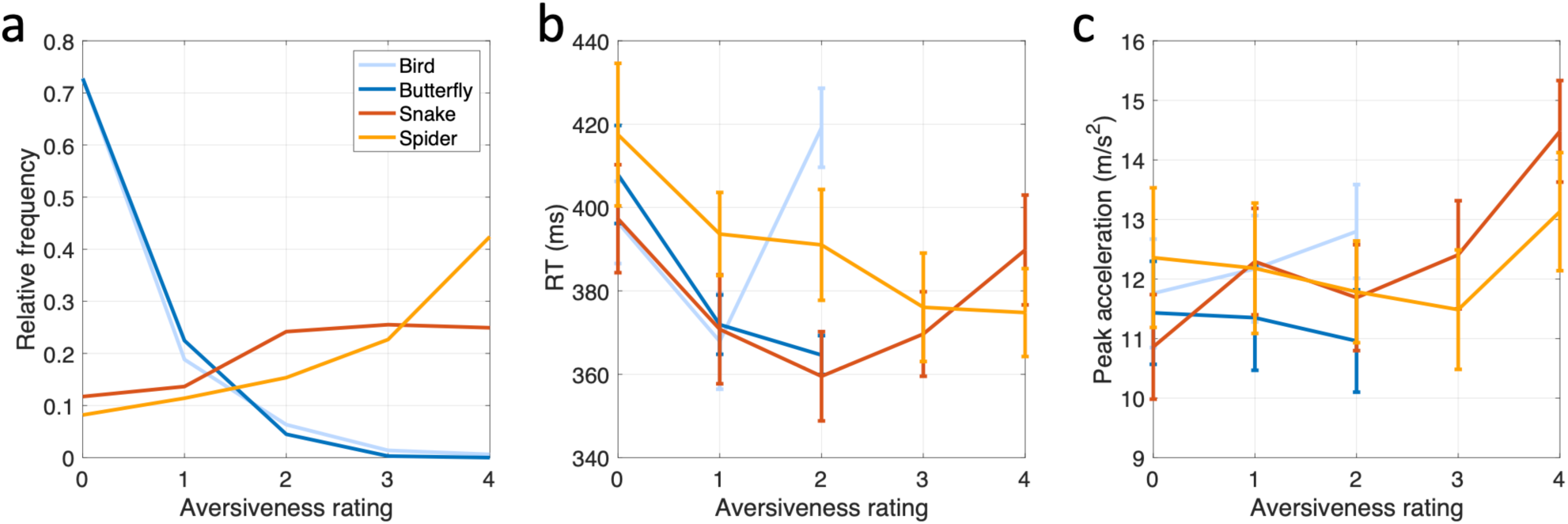
Association between reported aversiveness of the stimuli, and behavioral withdrawal responses. a) Relative frequency of different aversiveness ratings for the four animal categories. b) RT as a function of aversiveness rating. c) Peak acceleration as a function of aversiveness rating. Different line colors indicate different animal stimuli. The two lowest RT and peak acceleration values are not displayed for birds and butterflies because of small number of observations. Error bars display the SEM.

**Supplementary Table 1.** Correlation between fear of the stimulus and behavioral response across participants.

|  |  |  |  |  |  |  |
| --- | --- | --- | --- | --- | --- | --- |
| Name | Estimate | SE | t | p | Lower 95% CI | Upper 95% CI |
| **Reaction time (df = 224, R<sup>2</sup> adjusted = .87)** |  |  |  |  |  |  |
| (Intercept) | 331.58 | 5.02 | 66.07 | < .001 | 321.69 | 341.47 |
| Aversiveness rating | -1.58 | 2.46 | -0.64 | 0.52 | -6.42 | 3.26 |
| Bird | -6.20 | 4.19 | -1.48 | 0.14 | -14.47 | 2.06 |
| Snake | -1.95 | 6.25 | -0.31 | 0.76 | -14.28 | 10.37 |
| Spider | 8.00 | 7.09 | 1.13 | 0.26 | -5.98 | 21.98 |
| Condition_Withdraw | 71.90 | 8.11 | 8.86 | < .001 | 55.91 | 87.88 |
| Aversiveness:Withdraw | -8.86 | 2.22 | -4.00 | < .001 | -13.23 | -4.49 |
| **Peak acceleration (df = 224, R<sup>2</sup> adjusted = .98)** |  |  |  |  |  |  |
| (Intercept) | 7.47 | 0.58 | 12.95 | < .001 | 6.33 | 8.60 |
| Aversiveness rating | 0.00 | 0.07 | -0.02 | 0.98 | -0.14 | 0.14 |
| Bird | 0.05 | 0.11 | 0.48 | 0.63 | -0.16 | 0.26 |
| Snake | 0.39 | 0.17 | 2.32 | 0.02 | 0.06 | 0.71 |
| Spider | 0.35 | 0.19 | 1.85 | 0.07 | -0.02 | 0.72 |
| Condition_Withdraw | 4.35 | 0.77 | 5.66 | < .001 | 2.84 | 5.86 |
| Aversiveness:Withdraw | 0.14 | 0.06 | 2.34 | 0.02 | 0.02 | 0.25 |
| y ~ Animal + (Withdraw * Aversiveness) + (1 + Withdraw \| Participant) |  |  |  |  |  |  |

**Supplementary Figure 2.**
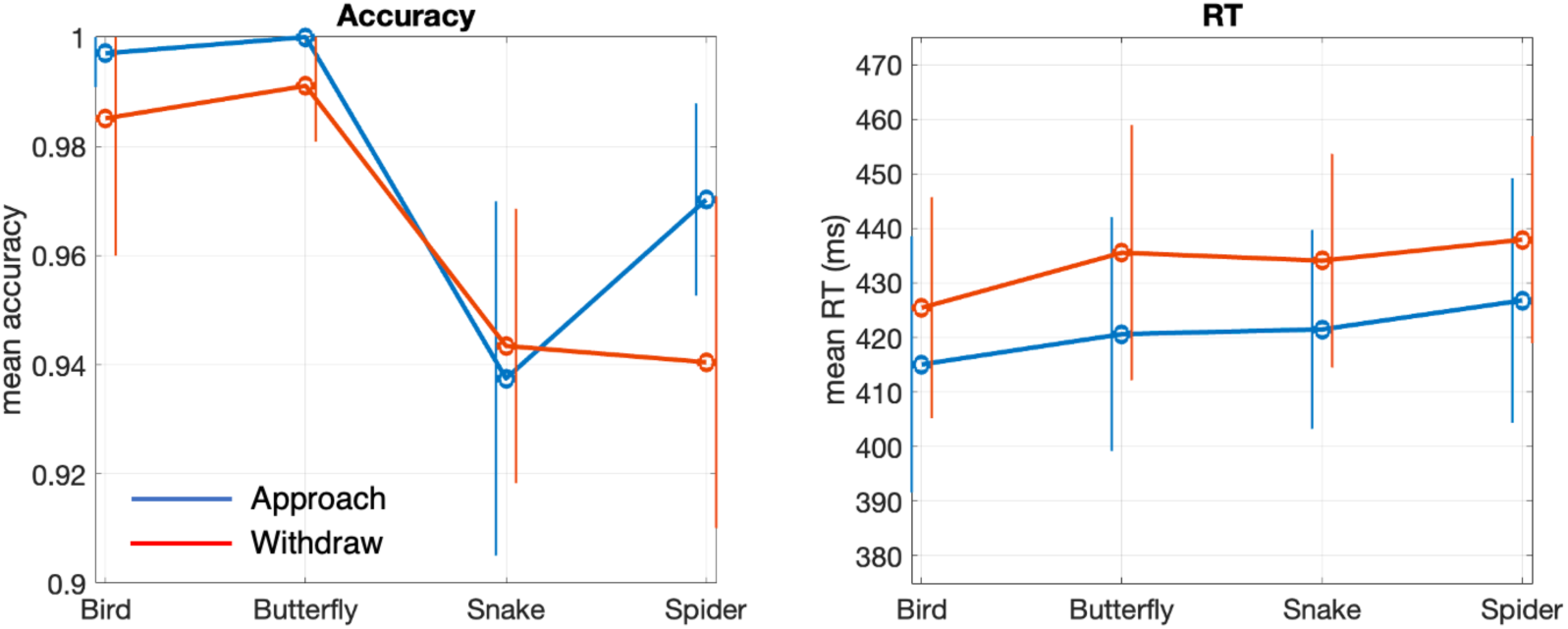
Results of the control experiment. Blue line is the button-press condition, and red line the button-release condition. Error bars display the 95% CI.

**Supplementary Table 2.** Results of mixed-effects regression analysis on RT in the control experiment (df = 2591, R^2^ adjusted = .34)

| Parameter | Estimate | SE | t | p | Lower 95% CI | Upper 95% CI |
| --- | --- | --- | --- | --- | --- | --- |
| Intercept (Butterfly, button-press) | 414.66 | 10.44 | 39.71 | < .001 | 394.19 | 435.14 |
| Bird | -1.34 | 6.46 | -0.21 | 0.84 | -14.00 | 11.32 |
| Snake | 4.91 | 6.46 | 0.76 | 0.45 | -7.77 | 17.58 |
| Spider | 9.29 | 6.46 | 1.44 | 0.15 | -3.37 | 21.96 |
| Button-release | 15.34 | 8.53 | 1.80 | 0.07 | -1.38 | 32.07 |
| Bird:Button-release | -11.12 | 7.75 | -1.43 | 0.15 | -26.33 | 4.08 |
| Snake:Button-release | -6.30 | 7.75 | -0.81 | 0.42 | -21.50 | 8.89 |
| Spider:Button-release | -4.54 | 7.74 | -0.59 | 0.56 | -19.71 | 10.64 |

$y \sim \text{Animal} * \text{ReleaseCondition} + (1 + \text{ReleaseCondition} | \text{Participant}) + (1 + \text{ReleaseCondition} | \text{stimulus})$

**Supplementary Table 3.** Results of the mixed-effects regression model on behavioral data of the EEG experiment.

| Parameter | Estimate | SE | t | p | Lower 95% CI | Upper 95% CI |
| --- | --- | --- | --- | --- | --- | --- |
| **Reaction time (df = 7903, $R^2$ adjusted = .30)** |  |  |  |  |  |  |
| Intercept (Butterfly) | 459.62 | 8.25 | 55.72 | < .001 | 443.45 | 475.79 |
| Snake | -12.37 | 4.29 | -2.88 | < .001 | -20.77 | -3.96 |
| Spider | -3.03 | 4.28 | -0.71 | 0.48 | -11.42 | 5.36 |
| Bird | -4.09 | 4.27 | -0.96 | 0.34 | -12.46 | 4.29 |
| Peak acceleration (z) | -4.05 | 1.12 | -3.63 | < .001 | -6.25 | -1.86 |
| **Peak acceleration (df = 7741, $R^2$ adjusted = .65)** |  |  |  |  |  |  |
| Intercept (Butterfly) | 10.23 | 0.66 | 15.46 | < .001 | 8.93 | 11.52 |
| Snake | 0.70 | 0.08 | 9.25 | < .001 | 0.55 | 0.85 |
| Spider | 0.54 | 0.08 | 7.17 | < .001 | 0.39 | 0.69 |
| Bird | -0.16 | 0.07 | -2.18 | 0.03 | -0.31 | -0.02 |
| RT (z) | -0.10 | 0.03 | -3.13 | < .001 | -0.16 | -0.04 |

$y \sim \text{Animal} + \text{RTorAcceleration} + (1 | \text{Participant}) + (1 | \text{stimulus})$

**Supplementary Figure 3.**
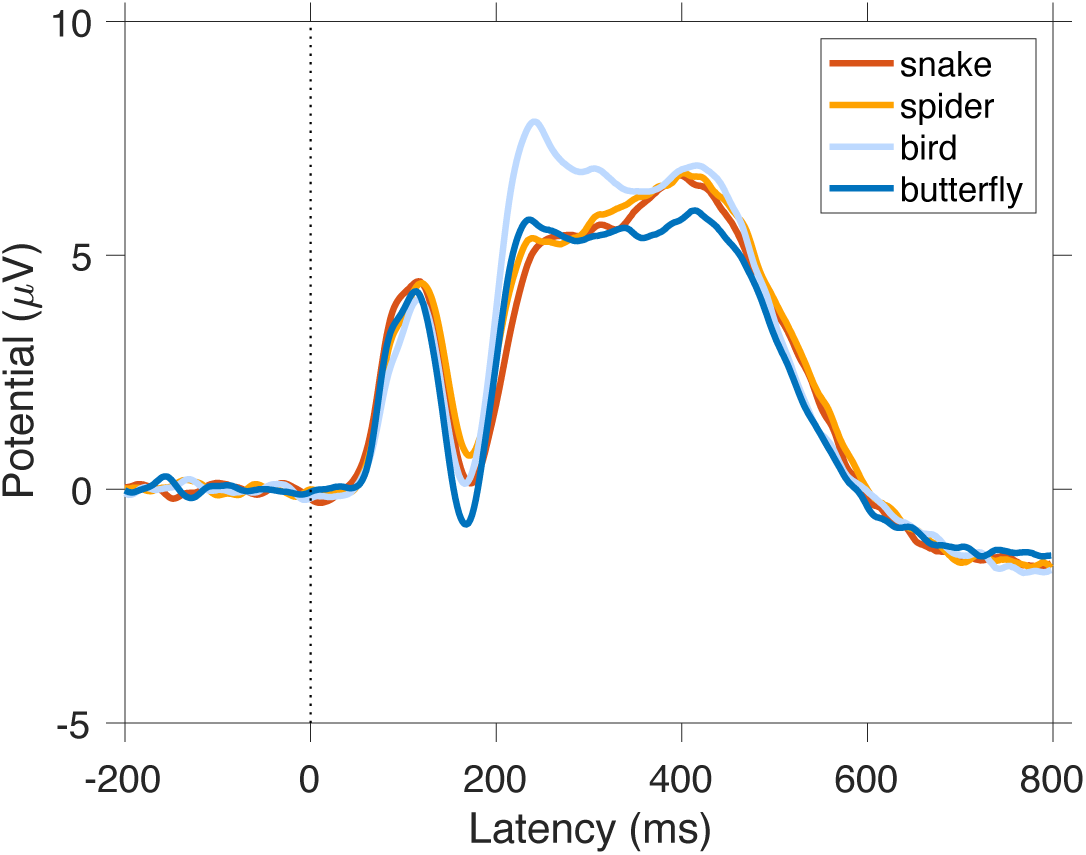
Grand average ERPs to different animal categories (mean of channels P7 and P8, i.e., electrode included in the N1 analysis).

**Supplementary Figure 4.**
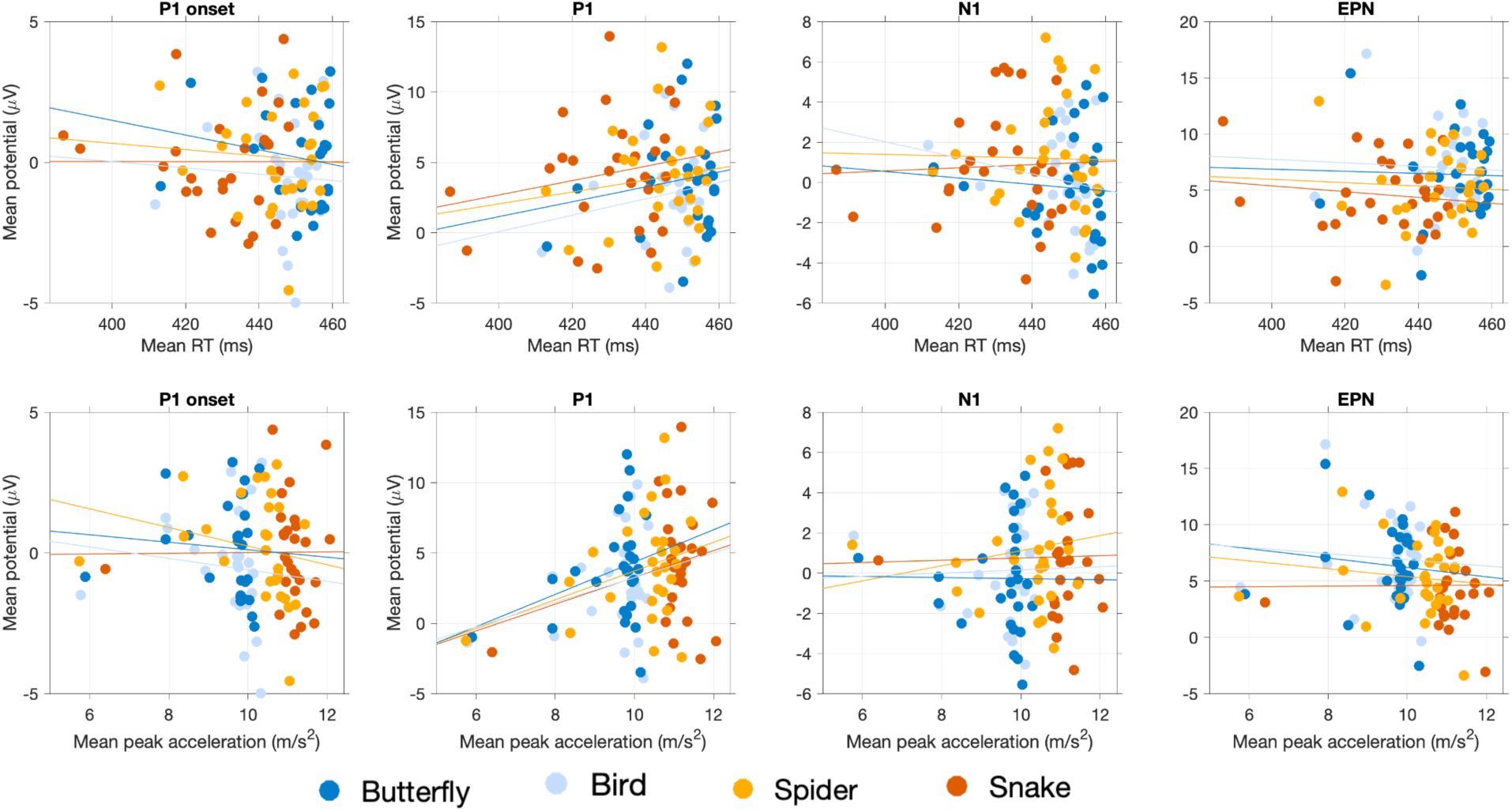
Between-participants association between behavioral metrics and ERPs. The top four plots display mean RT, and the bottom four plots the mean acceleration. The four columns of plots display relationship with different ERP waves: P1 onset, P1, N1, and EPN, respectively. Each dot represents one participant’s mean data. Different colors denote different animal stimulus categories. The lines are simple least-squares fits across the data points.

**Supplementary Figure 5.**
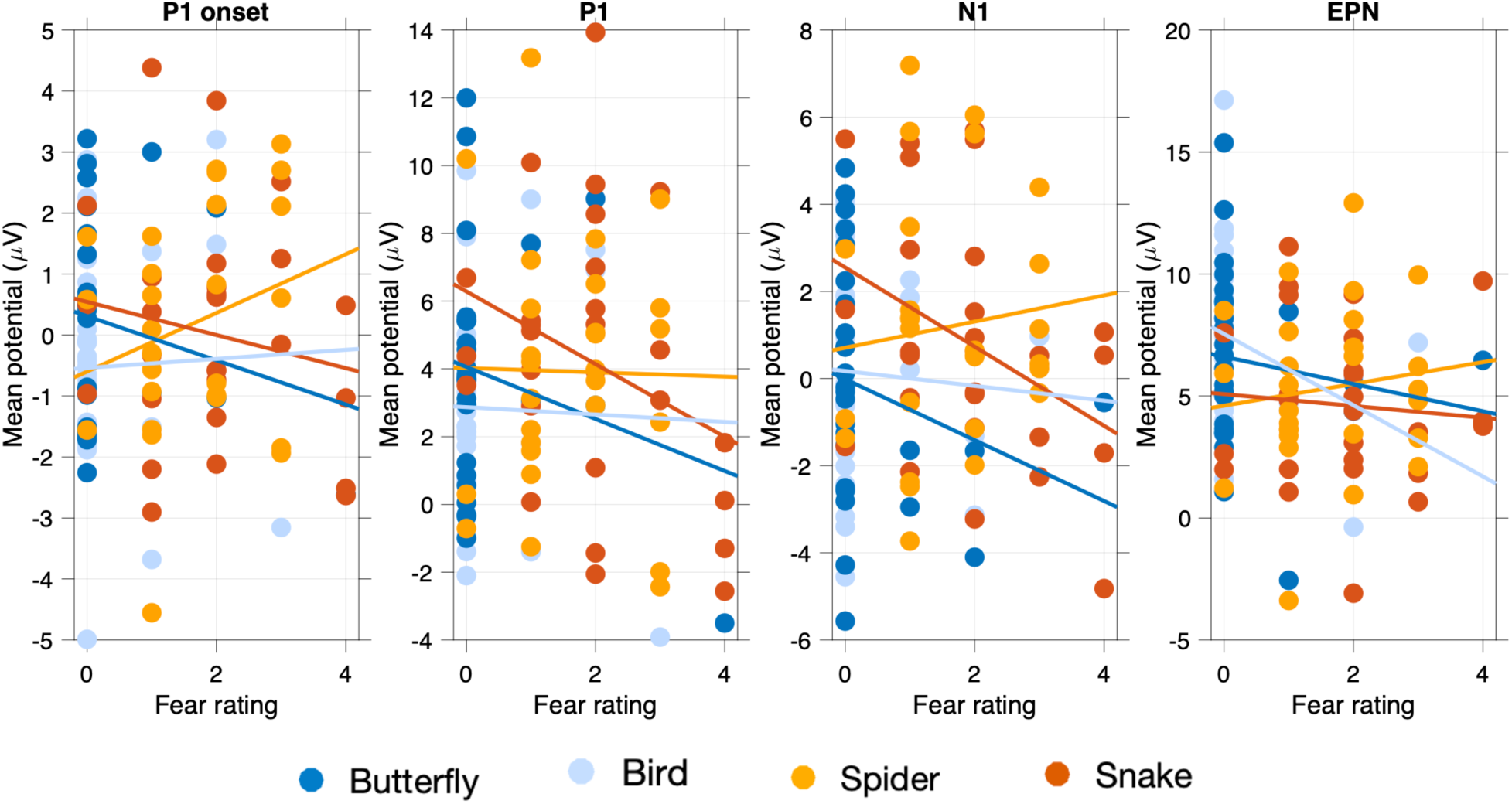
Between-participants association between fear rating and ERPs. The four columns of plots display relationship with different ERP waves: P1 onset, P1, N1, and EPN, respectively. Each dot represents one participant’s mean data. Different colors denote different animal stimulus categories. The lines are simple least-squares fits across the data points.

## Notes

https://osf.io/j6sqx/

### Competing Interest Statement

The authors have declared no competing interest.

### Summary of Updates

Revised writing, some additional analyses, and bit more extensive Discussion

